# Metabolic roles of uncultivated bacterioplankton lineages in the northern Gulf of Mexico “Dead Zone”

**DOI:** 10.1101/095471

**Authors:** J. Cameron Thrash, Kiley W. Seitz, Brett J. Baker, Ben Temperton, Lauren E. Gillies, Nancy N. Rabalais, Bernard Henrissat, Olivia U. Mason

## Abstract

Marine regions that have seasonal to long-term low dissolved oxygen (DO) concentrations, sometimes called ‘dead zones,’ are increasing in number and severity around the globe with deleterious effects on ecology and economics. One of the largest of these coastal dead zones occurs on the continental shelf of the northern Gulf of Mexico (nGOM), which results from eutrophication-enhanced bacterioplankton respiration and strong seasonal stratification. Previous research in this dead zone revealed the presence of multiple cosmopolitan bacterioplankton lineages that have eluded cultivation, and thus their metabolic roles in this ecosystem remain unknown. We used a coupled shotgun metagenomic and metatranscriptomic approach to determine the metabolic potential of Marine Group II Euryarchaeota, SAR406, and SAR202. We recovered multiple high-quality, nearly complete genomes from all three groups as well as those belonging to Candidate Phyla usually associated with anoxic environments-Parcubacteria (OD1) and Peregrinibacteria. Two additional groups with putative assignments to ACD39 and PAUC34f supplement the metabolic contributions by uncultivated taxa. Our results indicate active metabolism in all groups, including prevalent aerobic respiration, with concurrent expression of genes for nitrate reduction in SAR406 and SAR202, and dissimilatory nitrite reduction to ammonia and sulfur reduction by SAR406. We also report a variety of active heterotrophic carbon processing mechanisms, including degradation of complex carbohydrate compounds by SAR406, SAR202, ACD39, and PAUC34f. Together, these data help constrain the metabolic contributions from uncultivated groups in the nGOM during periods of low DO and suggest roles for these organisms in the breakdown of complex organic matter.

**Importance:** Dead zones receive their name primarily from the reduction of eukaryotic macrobiota (demersal fish, shrimp, etc.) that are also key coastal fisheries. Excess nutrients contributed from anthropogenic activity such as fertilizer runoff result in algal blooms and therefore ample new carbon for aerobic microbial metabolism. Combined with strong stratification, microbial respiration reduces oxygen in shelf bottom waters to levels unfit for many animals (termed hypoxia). The nGOM shelf remains one of the largest eutrophication-driven hypoxic zones in the world, yet despite its potential as a model study system, the microbial metabolisms underlying and resulting from this phenomenon—many of which occur in bacterioplankton from poorly understood lineages—have received only preliminary study. Our work details the metabolic potential and gene expression activity for uncultivated lineages across several low DO sites in the nGOM, improving our understanding of the active biogeochemical cycling mediated by these “microbial dark matter” taxa during hypoxia.

## Introduction

Hypoxia (dissolved oxygen [DO] below 2 mg·L^-1^/~62.5 μmol·kg^-1^) is dangerous or lethal to a wide variety of marine life, including organisms of economic importance (1). Hypoxia results from oxygen consumption by aerobic microbes combined with strong stratification that prevents reoxygeneation of bottom waters. These taxa are fueled primarily by autochthonous organic matter generated from phytoplankton responding to nitrogen input (1). Hypoxic zones have become more widespread globally through the proliferation of nitrogen-based fertilizers and the resulting increases in transport to coastal oceans via runoff (2). In the nGOM, nitrogen runoff from the Mississippi and Atchafalaya Rivers leads to bottom water hypoxia that can extend over 20,000 km^2^-one of the world’s largest seasonal “dead zones” (1). Action plans to mitigate nGOM hypoxia have stressed that increasing our “understanding of nutrient cycling and transformations” remains vital for plan implementation (3). These needs motivated our current study of the engines of hypoxic zone nutrient transformation: microorganisms.

Much of our current knowledge regarding microbial contributions to regions of low DO comes from numerous studies investigating naturally occurring, deep-water oxygen minimum zones (OMZs), such as those in the Eastern Tropical North and South Pacific, the Saanich Inlet, and the Arabian, Baltic, and Black Seas (4-11). In many of these systems, continual nutrient supply generates permanent or semi-permanent decreases in oxygen, sometimes to the point of complete anoxia (4). During these conditions, anaerobic metabolisms, such as nitrate and sulfate reduction and anaerobic ammonia oxidation, become prevalent (5, 9, 11-13). In contrast, nGOM hypoxia is distinguished by a seasonal pattern of formation, persistence, and dissolution (1); benthic contributions to bottom water oxygen consumption (14, 15); and a shallow shelf that places much of the water column within the euphotic zone (16). While parts of the nGOM hypoxic zone can become anoxic (1, 17), many areas maintain low oxygen concentrations even during peak hypoxia while the upper water column remains oxygenated (18-20).

The first studies of bacterioplankton assemblages during nGOM hypoxia showed nitrifying Thaumarchaea dominated (21) and could be highly active (22), suggesting a major role for these taxa in nGOM nitrogen cycling. However, many more poorly understood organisms from cosmopolitan, but still uncultivated “microbial dark matter” (23) lineages, such as Marine Group II Euryarchaeota (MGII), SAR406, and SAR202, also occurred in abundance (21, 22). While the likely functions of some of these groups have become clearer recently, all of them contain multiple sublineages that may have distinct metabolic roles. For example, the SAR202 lineage of Chloroflexi contains at least five subclades with distinct ecological profiles (24, 25), and the best understood examples have been examined in the context of complex carbon degradation in the deep ocean (25). Likewise, SAR406 represents a distinct phylum with numerous sublineages, and the bulk of metabolic inference comes from taxa in deep water OMZs (23, 26-28).

None of these groups have been studied in detail in shallow coastal waters, particularly in the context of seasonal hypoxia. Thus, we pursued a combined metagenomic/metatranscriptomic approach to i) elucidate the specific contributions of these uncultivated lineages to biogeochemical cycling in the nGOM during hypoxia, ii) evaluate the relative similarity of these organisms to their counterparts elsewhere, and iii) determine if other uncultivated lineages had eluded previous microbial characterization in the region due to confounding factors such as primer bias (29), 16S rRNA gene introns (30), or low abundance. Metagenomic binning recovered 20 genomes across seven uncultivated lineages including MGII, SAR406, and SAR202, and also from Candidate Phyla previously uncharacterized in the nGOM: Parcubacteria (23), Peregrinibacteria (31), and possibly PAUC34f (32) and ACD39 (33). Our results provide the first information on the likely potential function and activity of these taxa during hypoxia in the shallow nGOM and suggest novel roles for some of these groups that possibly reflect sublineage-specific adaptations.

## Results

### Study area

Our previous work used 16S rRNA gene amplicon data and qPCR to examine correlations between whole microbial communities, nutrients, and DO across the geographic range of the 2013 seasonal hypoxia (21). Here we selected six of those samples from offshore of the region between Atchafalaya Bay and Terrebonne Bay (D’, D, and E transects). These sites ranged considerably in DO concentration (~2.2 – 132 μmol·kg^-1^), and we chose them to facilitate a detailed investigation of the metabolic repertoire of individual taxa across the span of suboxic (1-20 μmol·kg^-1^ DO) to oxic (> 90 μmol·kg^-1^ DO) (5) water. Microbial samples from these sites were collected at the oxygen minimum near the bottom. Site depth ranged from 8 – 30 m, with the hypoxic (< 2 mg·L^-1^/62.5 μmol·kg^-1^) layer (at sites D2, D3, E2A, and E4) extending up to ~5 m off the bottom (Table S1).

### Metagenomic assembly yielded high quality genomes from multiple uncultivated lineages

Our initial assembly and binning efforts recovered 76 genomes. Using a concatenated ribosomal protein tree that included members of the Candidate Phylum Radiation (CPR) (34) (Fig. S1), CheckM (35) (Fig. S2), 16S rRNA genes and other single copy markers where available, and analyses of individual gene taxonomy (Fig. S3), we assigned 20 genomes to uncultivated “microbial dark matter” groups. These were six Marine Group II Euryarchaeota (MGII), five Marinimicrobia (SAR406), three in the SAR202 clade of Chloroflexi, and within Candidate Phyla (CP), one Parcubacteria (OD1), two Peregrinibacteria, and putatively, one ACD39 and two PAUC34f (Table 1, Supplemental Information). We further defined the MGII, SAR406, and SAR202 genomes into sublineages based on average amino acid identity (AAI), GC content, clade structure in the ribosomal protein tree, and 16S rRNA genes (Supplemental Information). SAR406 genomes belonged to two groups, A and B, corresponding to the previously established Arctic96B-7 and SHBH1141 16S rRNA gene clades (27). The three SAR202 genomes belonged to the previously established subclade I 16S rRNA gene clade (24). All genomes, with the exception of the Parcubacteria Bin 40, had estimated contamination of less than 6%, and in the majority of cases, less than 2%. Four of the six MGII genomes had estimated completeness (via CheckM) of greater than 61%, four of the five SAR406 greater than 73%, and all three SAR202 genomes were estimated to be greater than 83% complete. All CP lineages had at least one genome estimated to be greater than 71% complete (Table 1).

**Table 1.** Genome characteristics for the 20 bins associated with uncultivated lineages.

| IMG<br>Genome ID | Bin<br>Id | Taxonomy | Compl<br>% | Contam<br>% | Strain<br>het. | Scaff. | Longest<br>scaff.<br>(bp) | Size<br>(bp) | Genes | GC<br>(fract.) | Coding<br>density<br>(fract.) | Estim.<br>Compl.<br>Size<br>(Mbp) |
| --- | --- | --- | --- | --- | --- | --- | --- | --- | --- | --- | --- | --- |
| 2651870035 | 43-1 | Chloroflexi<br>(SAR202) | 90.3 | 0 | 0 | 69 | 161242 | 1972793 | 1882 | 0.52 | 0.88 | 2.2 |
| 2651870036 | 43-2 | Chloroflexi<br>(SAR202) | 88.6 | 4.1 | 15.4 | 134 | 157345 | 2402386 | 2373 | 0.52 | 0.91 | 2.7 |
| 2651870034 | 43 | Chloroflexi<br>(SAR202) | 83.2 | 0.1 | 100 | 179 | 90503 | 2475308 | 2392 | 0.53 | 0.89 | 3.0 |
| 2693429801 | 45 | Marinimicrobia<br>(SAR406) –B | 89.8 | 5.5 | 71.4 | 124 | 105140 | 2811623 | 2487 | 0.47 | 0.93 | 3.1 |
| 2651870052 | 45-1 | Marinimicrobia<br>(SAR406) –B | 85.2 | 0.1 | 100 | 144 | 100582 | 2410233 | 2235 | 0.49 | 0.93 | 2.8 |
| 2693429802 | 45-2 | Marinimicrobia<br>(SAR406) –B | 79.3 | 1.8 | 12.5 | 287 | 61408 | 2811444 | 2554 | 0.46 | 0.94 | 3.5 |
| 2651870053 | 51-1 | Marinimicrobia<br>(SAR406) –A | 73.6 | 1.7 | 50 | 75 | 191525 | 1901306 | 1835 | 0.39 | 0.95 | 2.6 |
| 2651870051 | 51 | Marinimicrobia<br>(SAR406) –A | 21.3 | 0 | 0 | 115 | 16923 | 578802 | 686 | 0.41 | 0.95 | 2.7 |
| 2651870038 | 15 | Euryarchaeota<br>(MGII) | 83.2 | 1.6 | 0 | 43 | 255599 | 1885130 | 1614 | 0.62 | 0.95 | 2.3 |
| 2651870039 | 17 | Euryarchaeota<br>(MGII) | 81.9 | 0.1 | 50 | 88 | 137319 | 1803861 | 1564 | 0.43 | 0.96 | 2.2 |
| 2651870037 | 14 | Euryarchaeota<br>(MGII) | 71.2 | 0.8 | 100 | 80 | 90069 | 1389909 | 1305 | 0.54 | 0.94 | 2.0 |
| 2651870040 | 18 | Euryarchaeota<br>(MGII) | 61.1 | 0.8 | 100 | 155 | 20633 | 1033226 | 1025 | 0.50 | 0.96 | 1.7 |
| 2651870042 | 38 | Euryarchaeota<br>(MGII) | 27.1 | 0 | 0 | 138 | 12893 | 615290 | 610 | 0.55 | 0.96 | 2.3 |
| 2651870041 | 17-1 | Euryarchaeota<br>(MGII) | 16.6 | 2.8 | 33.3 | 123 | 16952 | 538052 | 566 | 0.41 | 0.95 | 3.2 |
| 2693429807 | 13 | Unclassified<br>(ACD39) | 89.8 | 5.1 | 20 | 401 | 75104 | 4269849 | 3686 | 0.47 | 0.93 | 4.8 |
| 2693429799 | 40 | Parcubacteria<br>(OD1) | 71.9 | 15.1 | 75 | 97 | 65104 | 1086283 | 1208 | 0.52 | 0.92 | 1.5 |
| 2693429797 | 16 | Peregrinibacteria | 83.2 | 0.3 | 100 | 67 | 59308 | 1384712 | 1318 | 0.39 | 0.93 | 1.7 |
| 2693429798 | 39 | Peregrinibacteria | 49.9 | 0.3 | 100 | 131 | 18806 | 747520 | 809 | 0.45 | 0.95 | 1.5 |
| 2693429804 | 50 | Unclassified<br>(PAUC34f) | 84.8 | 1.4 | 0 | 455 | 76269 | 5346994 | 5484 | 0.58 | 0.92 | 6.3 |
| 2693429803 | 48 | Unclassified<br>(PAUC34f) | 51.9 | 2.2 | 0 | 470 | 20176 | 2566149 | 2596 | 0.55 | 0.94 | 4.9 |

### Unique roles for the ubiquitous MGII, SAR406, and SAR202 lineages in nGOM hypoxia

MGII comprised over 10% of the total community in some samples from 2013, and one MGII OTU also had a strong negative correlation with DO during 2013 hypoxia (21). Within our metagenomics dataset, MGII were more abundant in lower oxygen samples than in fully oxic ones, and the most abundant of the lineages reported here (Fig. S7). The majority encoded for aerobic, chemoheterotrophic metabolism, with no predicted genes for nitrogen or sulfur respiration except for a putative nitrite reductase (*nirK*) in a single genome-Bin 15 (Fig.1, Table S1). MGII genomic abundance correlated well with transcriptional abundance in most samples (Fig. 3), and we specifically found MGII cytochrome c oxidase expression throughout, though the levels and patterns differed depending on the gene and the source genome (Fig. 4, Table S1). Expression of the *nirK* gene occurred in the D2 and E2A samples-both suboxic. All but the most incomplete genome encoded for ammonia assimilation, making this a likely nitrogen source. Aggregate metabolic construction from multiple bins also indicated a complete TCA cycle, glycolysis via the pentose phosphate pathway, and gluconeogenesis (Fig. 1). Carbohydrate active enzyme (CAZy) genes can provide critical information on the relationships between microbes and possible carbon sources (36). We found few and these were largely restricted to glycosyltransferases (GT) in families 2 and 4, with activities related to cellular synthesis. In general, CAZy expression occurred for at least one gene in every genome and we detected expression of GT cellular synthesis genes in the E2A sample (Fig. 5), likely indicating actively growing cells.

**Figure 1.**
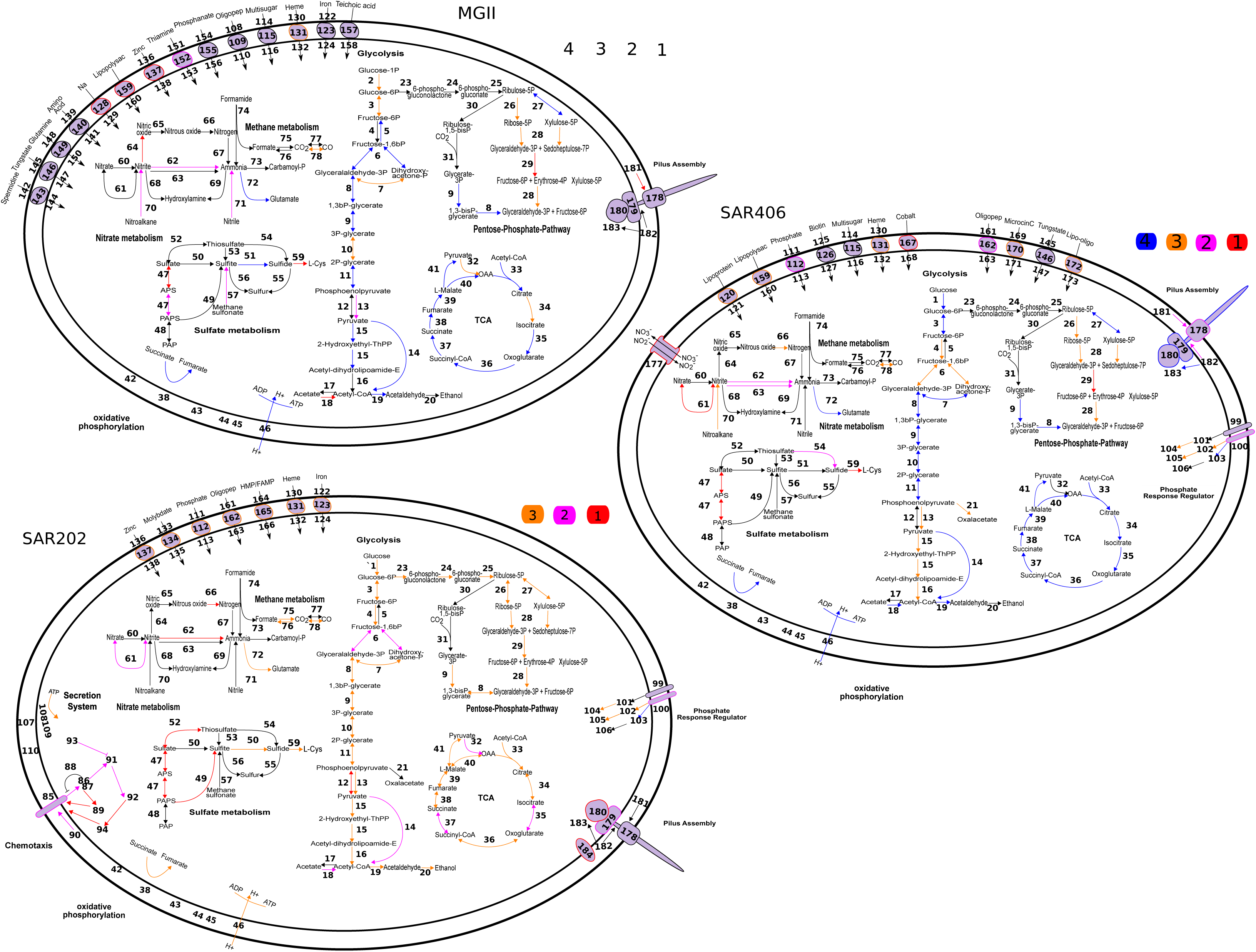
Metabolic reconstruction of Marine Group II Euryarchaeota, SAR406, and SAR202, based on the top three or four most complete genomes. Colors indicate pathway elements based on the number of genomes in which they were recovered, according to the key. Black outlines and/or arrows indicate genes that were not observed. Numbers correspond to annotations supplied in Table S1.

SAR406 represented over 5% of the population in some locations during hypoxia in 2013, and one abundant OTU was negatively correlated with DO (21). Metagenomic read recruitment to the SAR406 bins confirmed this trend, with greater recruitment in the suboxic samples relative to dysoxic or oxic (Fig. S7). Total RNA recruitment was strongest to Bins 45 and 51-1, though most bins showed a RNA to DNA recruitment ratio > 1 in at least one sample, indicating these taxa were likely active (Fig. 3). Despite their affinity for low oxygen environments, the SAR406 genomes encoded a predicted capacity for aerobic respiration (Fig. 1), and we found expression of cytochrome c oxidases in even the lowest oxygen samples (Fig. 4). The Group B genomes encoded both high and low-affinity cytochrome c oxidases (37), whereas the high affinity (cbb3-type) oxidases were not recovered in the Group A genomes (Table S1), which may indicate sublineage-specific optimization for different oxygen regimes.

Sublineage variation also appeared in genes for the nitrogen and sulfur cycles. Group B genomes all contained predicted nitrous oxide reductases *(nosZ)* and *nrfAH* genes for dissimilatory nitrite reduction to ammonium (defined here as DNRA, although this acronym frequently refers to nitrate, even though that is a misnomer (38)). The *nrfA* genes formed a monophyletic group with *Anaeromyxobacter dehalogenans* 2CP-1, an organism with demonstrated DNRA activity (38) (Fig. S8A). The genes also contained conserved motifs diagnostic of the *nrfA* gene (38) (Fig. S8B, C). We observed expression of *nrfAH* and *nosZ* at the sites with the lowest DO concentrations (D2, E2A, and E4) and expression appeared to have a negative relationship with DO concentration (Fig. 4). The Bin 51-1 Group A genome contained predicted *narHI* genes for dissimilatory nitrate reduction, which we did not find in the Group B genomes. We observed expression of SAR406 *narHI* only in the lowest DO sample from station E2A (Fig. 4). Two Group B SAR406 genomes had predicted *phsA* genes for thiosulfate reduction to sulfide (and/or polysulfide reduction (39)), as previously described from fosmid sequences (27). We detected transcripts for these genes only in samples E2A and E4, the two lowest DO samples (Fig. 4). Many of the anaerobic respiratory genes were co-expressed with cytochrome c oxidases, indicating a potential for either co-reduction of these alternative terminal electron acceptors or poising of these organisms for rapid switching between aerobic and anaerobic metabolism (40).

All SAR406 genomes had numerous genes for heterotrophy. We found CAZy genes in all major categories except polysaccharide lyases, and expression for most of these genes in both Group A and Group B genomes in one or more samples (Fig. 5). Notable carbohydrate compounds for which degradation capacity was predicted include cellulose (glycoside hydrolase (GH) families GH3, GH5; carbohydrate binding module (CBM) family CBM6), starch (GH13), agar and other sulfated galactans (GH2, GH16), chitin (GH18), xylan (GH30, CBM9), and peptidoglycan (GH23, GH103, CBM50). The genomes contained putative transporters for a variety of dissolved organic matter (DOM) components including nucleosides, amino and fatty acids, and oligopeptides (Table S1). We also found numerous outer membrane transporters, including cation symporters; Outer Membrane Receptors (OMR-TonB-dependent), which play important roles in transport of metals, vitamins, colicins, and other compounds; Outer Membrane Factors (OMF); and most genomes also had large numbers of duplicated genes (24 in Bin 45-2), identified via hidden Markov model searches against the SFam database (41), annotated as “Por secretion system C-terminal sorting domain-containing protein,” some of which were associated with GH16. These genes likely play a role in sorting C-terminal tags of proteins targeted for secretion via the Por system, which is essential for gliding motility and chitinase secretion in some Bacteroidetes (42). The extensive gene duplication may indicate expanded and/or specialized sorting functionality, and suggests an emphasis on protein secretion in this group. Expression of a membrane-bound lytic murein transglycosylase D (GH23) involved in membrane remodeling also supports the idea of active and growing cells from Group A in all samples (Table S1).

We detected Chloroflexi 16S rRNA gene sequences during 2013 hypoxia at up to 5% of the community (21), and recovered three mostly complete SAR202 Chloroflexi genomes in this work. Although present at lower abundance than MGII and SAR406 (Fig. S7), these genomes showed relatively high activity in some samples (Fig. 3). Like subclade III and V, subclade I organisms likely respire oxygen. However, we also found *napAB* and *nosZ* genes for nitrate and nitrous oxide reduction, respectively (Fig. 1). As in SAR406, we detected concurrent expression of these genes with cytochrome c oxidases in the lowest DO samples (Fig. 4) (Fig. 4, Table S1).

The SAR202 genomes have numerous transporters, many with predicted roles in organic matter transport, which supports previous observations of DOM uptake (43). In particular, SAR202 genomes had considerably more Major Facilitator Superfamily (MFS) transporters than the other genomes in this study (Table S1) and those of the subclade III genomes (25), and SFam searches revealed a the majority of these shared annotation as a “Predicted arabinose efflux permease” (SFam 346742). MFS genes transport numerous diverse substrates, such as sugars and amino acids, through coupling with an ion gradient, and can be associated with either uptake or export of compounds (44). SAR202 genomes also had between 53 and 66 predicted ABC transporters.

The SA202 genomes encoded a number of duplicated genes in specific gene families. The largest gene family expansion that we observed was associated with SFam 6706, with between 46 and 48 genes in this family encoded in each genome. Most of these (121/142) were annotated as either a “galactonate dehydratase” or a “L-alanine-DL-glutamate epimerase.” Galactonate dehydratase catalyzes the first step of the pathway to utilize D-galactonate in central carbon metabolism via the pentose phosphate pathway. The large number of genes in these categories likely indicates some divergence for alternative roles as this group belongs broadly to the COG4948 “L-alanine-DL-glutamate epimerase or related enzyme of enolase superfamily.”

All genomes also had numerous dehydrogenases as reported for the subclade III genomes (25). Specifically, SFams 346640 and 1639 were the third and fourth most abundant, with 16-18 and 13-15 genes in each family, respectively, across the three genomes. Genes in these families were annotated as “short-chain alcohol dehydrogenase family,” “3-alpha (or 20-beta)-hydroxysteroid dehydrogenase,” “meso-butanediol dehydrogenase,” and others. These match the annotations of the subclade III genomes, and suggest a similar role in conversion of alcohols to ketones (25). The SAR202 genomes have comparatively few CAZy genes relative to the other genomes. GH15 and GH63 suggest starch degradation, and GH105 pectin degradation, and we detected expression of multiple genes in these categories across samples (Fig. 5, Table S1).

### Other Candidate Phylum organisms in nGOM hypoxia

In contrast to the abundant and cosmopolitan MGII, SAR406, and SAR202 clades, we also recovered genomes from several groups that were either previously undetected in the nGOM or very rare. Although these taxa likely do not contribute the biomass of more populous clades, their genomes provide important insight into their functional potential during hypoxia. The Bin 13 genome (possibly ACD39) also had the highest relative activity compared to all the other genomes in our study (Fig. 3), underlining the point that low abundance does not automatically equate to low metabolic impact. Bin 13 had predicted aerobic respiration with both high and low-affinity cytochrome c oxidases (Fig. 2). The low affinity oxidases contributed more reads in the samples where we could detect expression (Table S1). The genome contained numerous predicted CAZy genes in the glycosyltransferase and glycoside hydrolase categories, spread across multiple families in each (Table S1). Notable degradation capacity included starch (GH13) and peptidoglycan (GH23, GH103, GH104).

**Figure 2.**
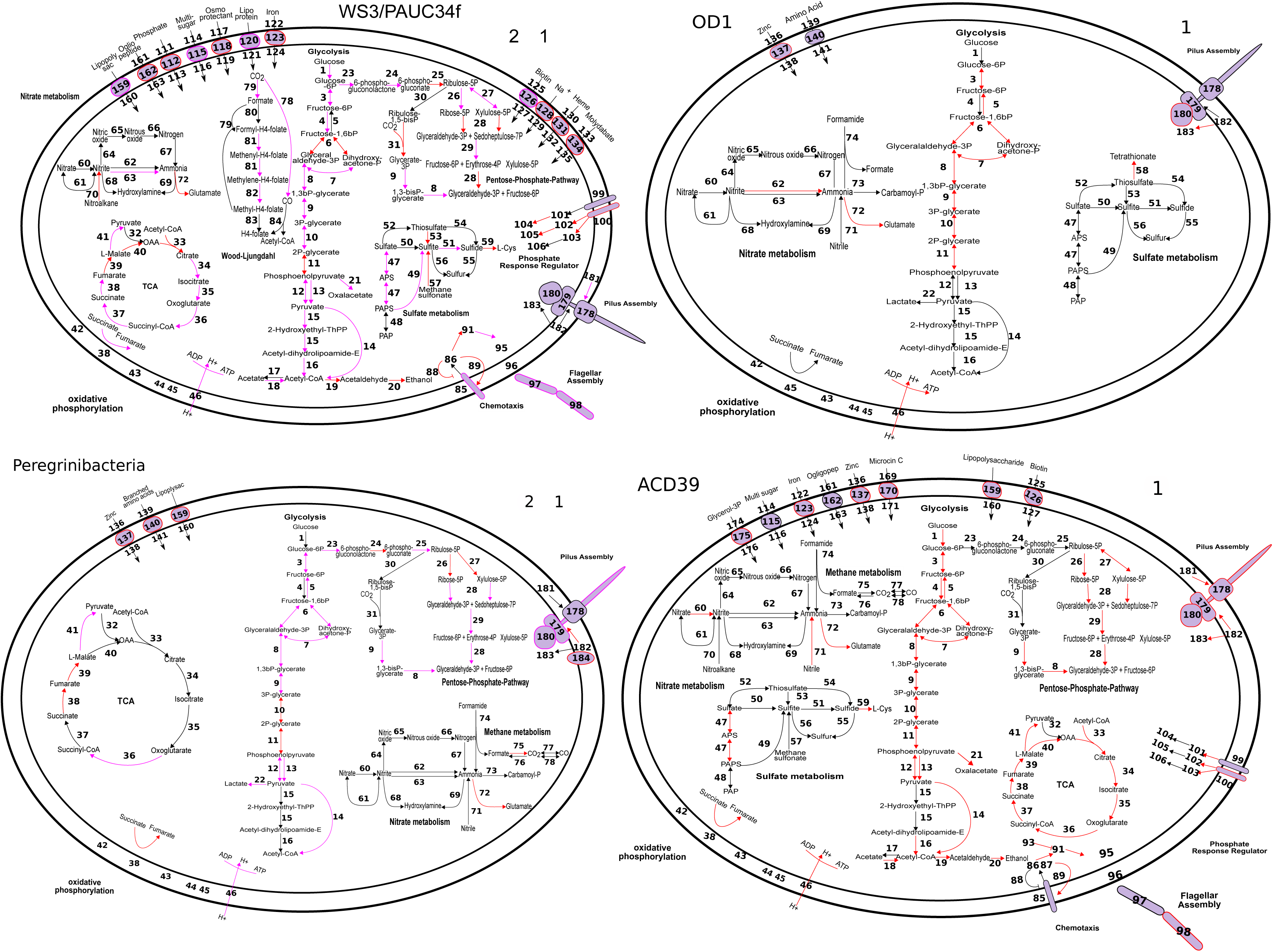
Metabolic reconstruction of the Candidate Phylum members PAUC34f, Parcubacteria (OD1), Peregrinibacteria, and ACD39 (Bin 13). Colors indicate pathway elements based on the number of genomes in which they were recovered, according to the key. Black outlines and/or arrows indicate genes that were not observed. Numbers correspond to annotations supplied in Table S1.

Bin 13 had ~ 80 ABC transporter genes, and similarly to the SAR406 genomes, numerous outer membrane transporters, including the OMR and OMF families. We predict complete glycolysis/gluconeogenesis pathways and a TCA cycle. We recovered paralogous pilus subunit genes, chemotaxis genes, and a partial flagellar assembly. Furthermore we detected relatively high expression of the *flgLN* flagellin genes in samples D2, D3, E2A, and E4 (Table S1) suggesting active motility in these environments. Several other Bin 13 genes were among the most highly expressed in all samples, but could only be classified as hypothetical (Table S1). Similarly, the three most populous SFams in Bin 13, according to number of genes (n=16, 15, and 13) also linked to genes annotated as hypothetical proteins with either tetratricopeptide, HEAT, TPR, or Sel1 repeats. Although currently obscure, these and the highly expressed hypothetical genes represent important targets for future research into the function of this group.

Bins 50 and 48 were lower in abundance than SAR202 genomes (Fig. S7, PAUC34f), with no observable trend associated with oxygen levels (Fig. S7). These genomes encoded flagellar motility, aerobic respiration, glycolysis via the pentose-phosphate pathway, gluconeogenesis, assimilatory sulfate reduction, and DNRA (Fig. 2). The *nrfA* subunit from both genomes grouped in the same monophyletic clade as those from SAR406 (Fig. S8A), and had similar conserved motifs (Fig. S8B, C). However, we note that the *nrfAH* gene sets for Bins 50 and 48 occurred on relatively short contigs (5650 and 5890 bp, respectively), so the metabolic assignment cannot be corroborated as definitively as that for SAR406. The Bin 50 genome was among the more active in our analysis (Fig. 3), and we detected highest expression of cytochrome c oxidase components in samples E2A and E4 (Fig. 4). DNRA gene expression was low but observable in the same samples. We also recovered a partial gene for the ribulose-bisphosphate carboxylase (RuBisCO) large subunit, but this fragment was on a very short contig (3954 bp), and we did not detect expression in any of our samples, so we cannot rule out that this gene occurred on a contaminating contig.

**Figure 3.**
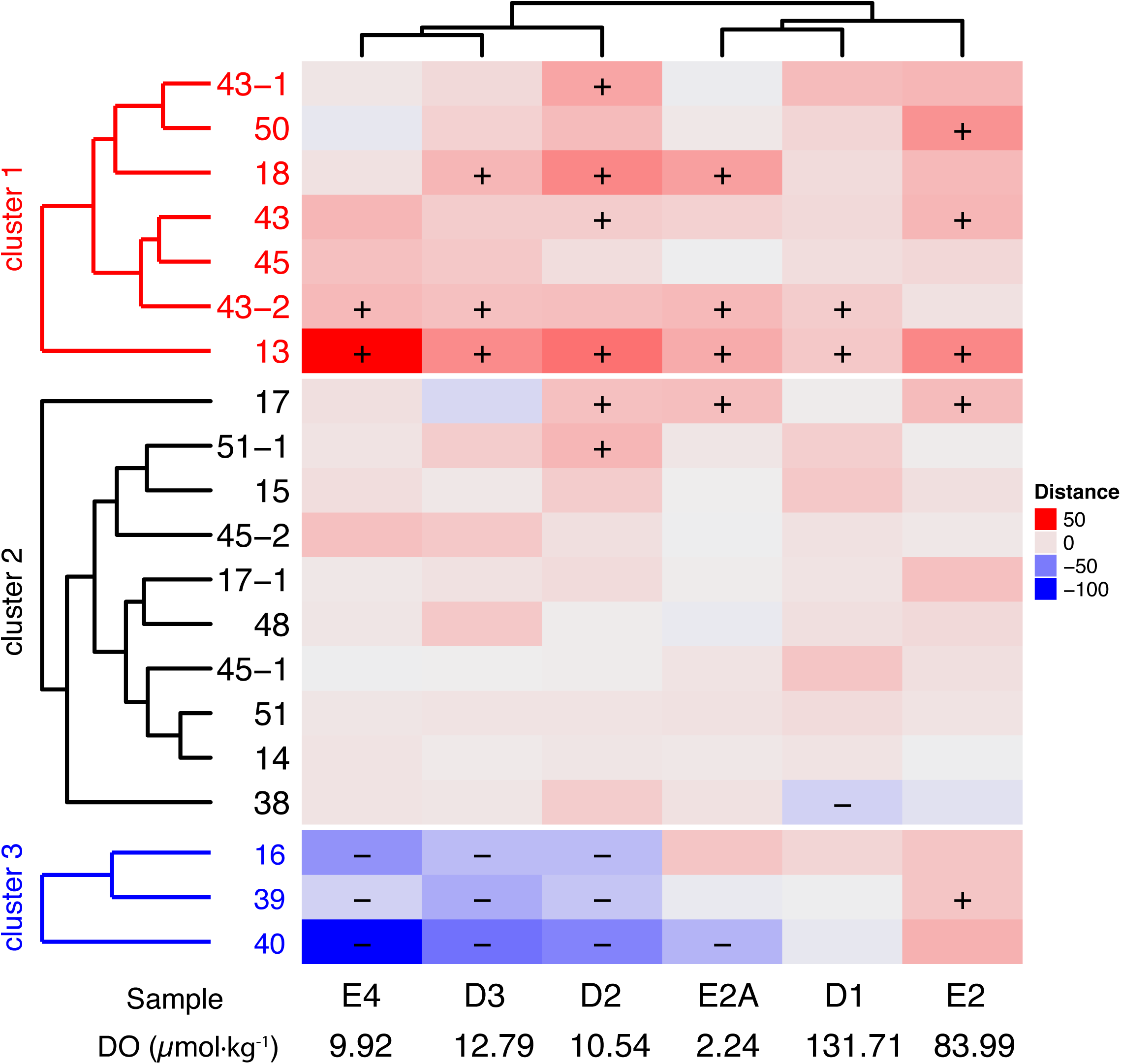
Relative DNA to RNA recruitment rank for each genome, by sample. Colors indicate the relative difference in the ratio of rank based on total RNA and DNA mapping. Red indicates a higher RNA recruitment rank compared to DNA recruitment rank, and vice-versa for blue. + and – symbols indicate bins where the rank-residual from the identity in RNA vs. DNA read mapping was more or less than one standard deviation beyond 0, respectively. Dendrograms were calculated using an Unweighted Pair Group Method with Arithmetic Mean (UPGMA) from Euclidian distances of rank residuals across all samples and bins.

**Figure 4.**
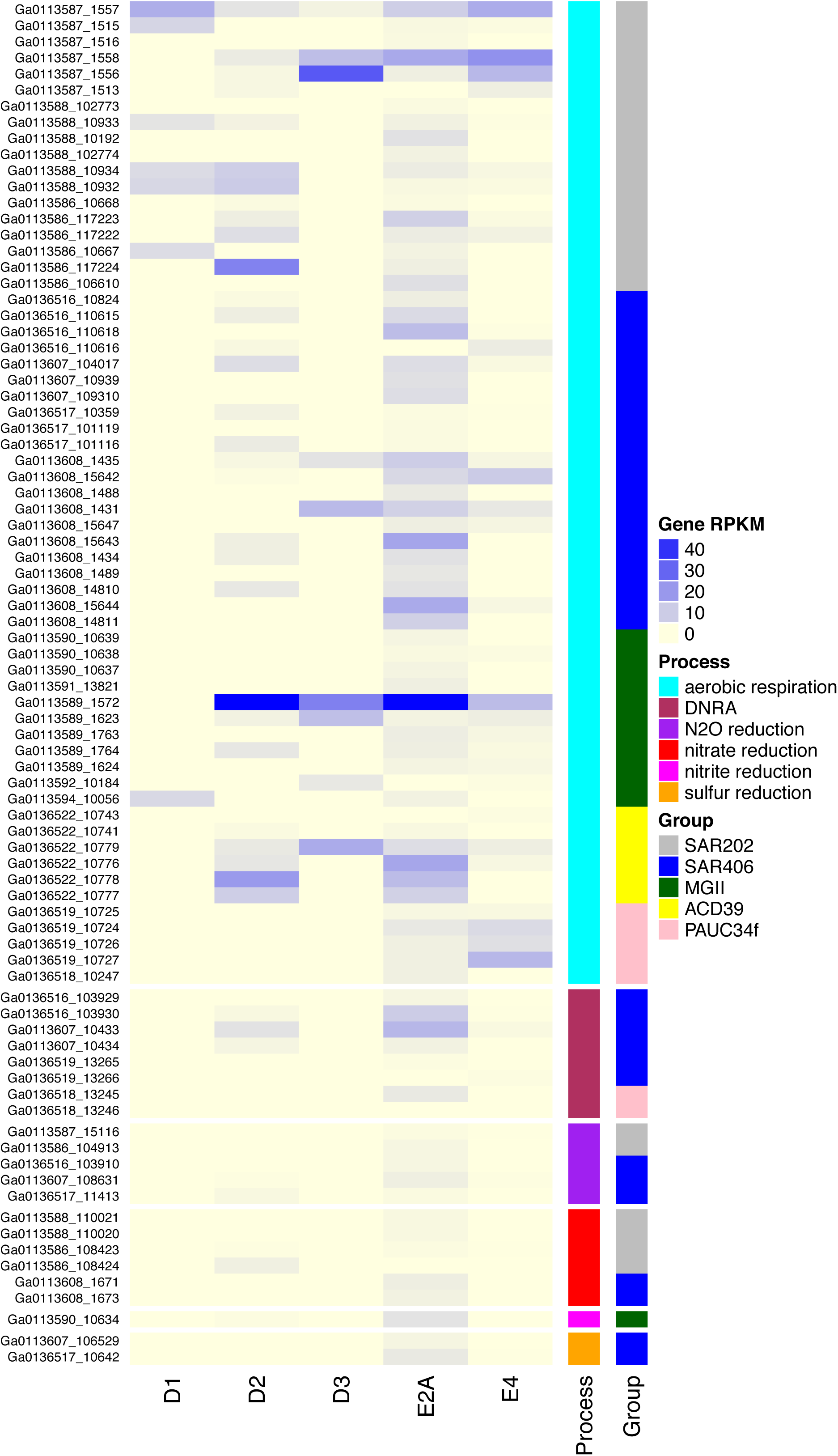
Expression of predicted respiratory genes. RPKM values of RNA recruitment for each gene, by sample, are depicted with colors according to the key (yellow to blue follows increasing intensity). Genes are grouped by bin, taxonomic affiliation, and specific respiratory process. DNRA-dissimilatory nitrite reduction to ammonia.

**Figure 5.**
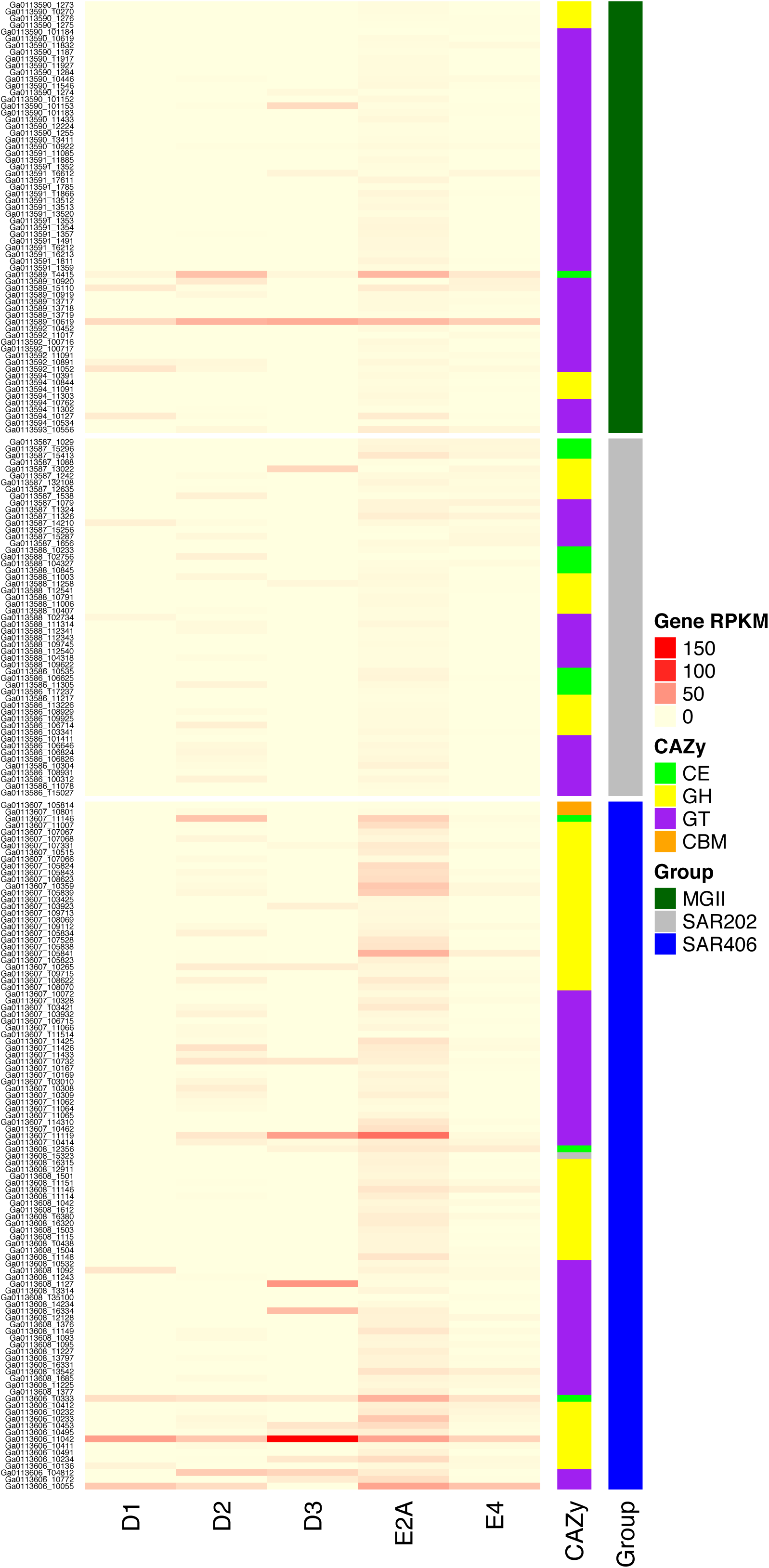
Expression of predicted CAZy genes. RPKM values or RNA recruitment for each gene, by sample, are depicted with colors according to the key (yellow to red follows increasing intensity). Genes are grouped by bin, taxonomic affiliation, and general CAZy categories. CE-carbohydrate esterase; GH-glycoside hydrolase; GT-glycosyltransferase; CBM-carbohydrate binding module.

The Bin 50 and 48 genomes had abundant CAZy genes in all categories, suggesting a highly flexible metabolic repertoire for carbon acquisition. They contain possible capacity for breakdown of starch (GH13, CBM48), peptidoglycan (GH23, CBM50), fructose-based oligosaccharides (GH32), and hemicellulose (GH2, GH3, GH43). Notably these genomes were the only ones with predicted polysaccharide lyases (PL) among those compared (with the exception of a single predicted PL gene in SAR406-Table S1). PL genes cleave uronic-acid containing polysaccharides (45). These organisms seem particularly adapted for pectin (PL1, PL2, PL9, PL10, PL11, PL22, GH78) and alginate (PL15, PL17) degradation-both compounds are common cell wall components of green and brown algae, respectively.

In line with the algal cell wall degradation ability, we detected a large expansion (102 genes in Bin 50) of sulfatase genes in SFam 1534, annotated predominantly as either “arylsulfatase A” or “choline-sulfatase.” Arylsulfatases cleave sulfate esters, usually to supply microbes with a source of sulfur, and can be located intracellularly or in membranes (46).

Choline-sulfatases cleave choline sulfate to choline and sulfate, with downstream use for the former as a carbon source or osmoprotectant and the latter as a sulfur source (47). Given the predicted assimilatory sulfate reduction pathway in Bins 50 and 48, this is a logical means to obtain sulfur for the group. We observed large expansions in galactonate and other dehydratases (as in SAR202, above-SFam 6706 n=42 in Bin 50), as well as numerous ABC transporter permeases (SFam 4442), which match the transporter predictions via IMG: 117 predicted genes for ABC transporters in all. These genomes also had numerous OMF and OMR transporter genes (Table S1). The large number of transporters and protein family expansions correspond to the relatively large expected genome sizes (between 5 and 6 Mbp).

We also recovered genomes associated with CPR taxa usually associated with anoxic environments: two Peregrinibacteria and one from the Uhrbacteria subclade of the Parcubacteria (formerly OD1). All three genomes could be assigned taxonomically with high confidence based on their positions in the ribosomal protein tree (Fig. S1) and via gene annotations (Fig. S3). We note that although the Peregrinibacteria bins (16 and 39) had very low predicted contamination, the Parcubacteria Bin 40 has 15% predicted contamination (75% of which we attribute to strain heterogeneity in the bin) (Table 1). Recovery of Parcubacteria from a coastal marine system is unusual, but not unprecedented. Parcubacteria single-cell genomes have been identified in marine and brackish sources (23), and we previously identified 26 rare OTUs assigned to the phylum in nGOM hypoxia (21). That number of OTUs may explain why we observed 20 single copy marker genes present in two copies in Bin 40 (Table S1).

In contrast to Parcubacteria, Peregrinibacteria have thus far only been found in terrestrial subsurface aquifers (31, 33, 48, 49), and remained undetected in our amplicon survey (21). Both groups occurred in low relative abundance to the other taxa in this study, and showed the lowest activity (Fig. 3). Consistent with previous reports of obligate fermentative metabolism by Parcubacteria and Peregrinibacteria (23, 30, 31, 48), we identified no respiratory pathways for these taxa (Fig. 2) and they trended towards greater abundances in the lowest DO samples (Fig. S7). In spite of relatively high predicted genome completion, we found very few CAZy genes, and those were mostly restricted to glycosyltransferases (Table S1) probably involved in capsular polysaccharide synthesis. While these organisms had low relative abundance to the other groups (Fig. S7), we did observe activity in some samples (Fig. 3-E2, E2A, D1).

## Discussion

This work provides the first reconstruction of multiple nearly complete genomes from uncultivated bacterioplankton during nGOM hypoxia. Although we define roles for MGII, SAR406, SAR202, Bin 13 and Bins 50/48 as aerobic heterotrophs, we also observed concurrent expression of genes associated with anaerobic metabolism in SAR406 (nitrate reduction, DNRA, nitrous oxide reduction, and sulfur reduction), SAR202 (nitrate and nitrous oxide reduction), MGII (nitrite reduction), and Bins 50/48 (DNRA) in suboxic samples with the lowest measured DO concentrations. Simultaneous utilization of multiple electron acceptors with different redox potentials likely indicates an abundant supply of electron donors (50), may denote niche partitioning within group sublineages at a finer level of taxonomic resolution than we observed, or indicate poising of taxa for rapidly changing chemical gradients (40). An organism’s set of CAZy genes often gives insights into its biology, in particular into nutrient sensing and acquisition. All taxa examined in this study had predicted chemoorganoheterotrophic metabolism, and the CAZy genes found in these genomes suggest that SAR406, SAR202, Bin 13, and Bins 50/48 participate in the degradation of complex organic matter resulting from the detritus of larger organisms. This matches the general model of hypoxic zone oxygen consumption resulting from sinking organic matter provided by algal blooms in surface waters (1). The observed activity of obligate fermentative groups Parcubacteria and Peregrinibacteria also suggests that anoxic pockets occur in the water column where these organisms can thrive.

Marine group II (MGII) is a broadly distributed archaeal clade, with members found in different marine (51, 52), and sedimentary (53), environments. Previous work during 2012 and 2013 hypoxia indicated a proliferation of archaeal taxa in both the Thaumarchaea and MGII phyla (21, 22). The prevalence of MGII among lower oxygen samples in the hypoxic zone is somewhat surprising, considering that they are commonly associated with aerobic environments (52). However, oxygen was still present in even the lowest DO samples (Fig. 3), and MGII success likely had more to do with the carbon content than oxygen levels. These nGOM MGII appear to be metabolically similar to those described in previous work: MGII have been shown to be dominant in water column environments associated with blooms in productivity, for example at deep-sea hydrothermal plumes (51). Thus, the increased availability of organic matter (proteins and carbohydrates), thought to be preferred substrates for MGII (54, 55), probably explains their abundance.

Another cosmopolitan group found in our samples was SAR406 or Marine Group A. These organisms were discovered over 20 years ago (28, 56), and the clade has recently been proposed as the phylum “Marinimicrobia” (23). SAR406 occur in numerous marine (5, 23, 26, 28, 57), sedimentary (23), and even oil reservoir (58) environments. They are prevalent in deeper ocean waters (28, 57, 59) and prefer lower oxygen concentrations in OMZs (5, 26, 60). Our genomes had larger estimated genome sizes-2.6-2.7 Mbp (Group A) and 2.8-3.5 Mbp (Group B)-compared to 1.1-2.4 Mbp from single-cell genomes (23). Overall GC content, however, was in the range of the 30-48% reported for fosmids (27) and single-cell genomes (23). The lower GC Group A genomes specifically had a similar GC content to the Arctic96B-7 fosmids, matching their predicted phylogenetic affiliation (see below) (27).

Our data now also define roles for them in the eutrophication-driven hypoxia of the nGOM. Previous metabolic reconstructions of SAR406 predicted aerobic metabolism (23) and sulfur reduction (27), which our data confirm, although the sulfur reduction genes were only found in Group B organisms (Table S1). Our genomes also suggest multiple nitrogen cycling roles that appear to be organized by sublineages within the phylum, and sublineage specific presence of both high and low affinity cytochrome c oxidases. The Group B organisms group with the early diverging SBH1141 clade (27), for which no previous genome data exist. Group B organisms contained both types of cytochrome c oxidases, *nosZ* and *nrfAH* genes, whereas Group A organisms, sister to the Arctic96B-7 clade, contained the low affinity cytochrome c oxidases only, and additionally *narHI* genes not found in Group B. The unique roles predicted for these taxa are not surprising given the diversity of the SAR406 clade and the genetic distances between Group A and B (Fig. S4). The fosmids associated with the Arctic69B-7 clade contained genes for oxidative stress and sulfur reduction (27), although we only found sulfur reduction genes in the distantly related Group B genomes. The ArcticB96-7 clade may be diverse enough to encompass differing metabolic strategies, but the variable presence of *phsA* genes in this group may simply be due to incomplete genomic data. In addition to sublineage-specific respiratory characteristics, our results also generate specific hypotheses about organic matter metabolism in SAR406: likely degradation capacity for cellulose, starch, agar, xylan, and peptidoglycan; transport of nucleosides, amino and fatty acids, and oligopeptides; and substantial gene duplication associated with protein secretion for possible extracellular metabolism.

Together these data suggest that during nGOM hypoxia SAR406 members degrade complex carbohydrates fueled by aerobic respiration, and supplemented with facultative anaerobic respiration of nitrate, nitrite, or sulfur compounds.

Members of the SAR202 clade of Chloroflexi also inhabit a wide variety of marine environments (24), frequently in deeper waters (24, 43, 57, 59, 61) and remain functionally understudied because genome data for SAR202 have been lacking. Landry and colleagues recently described the properties for several single-cell genomes representing SAR202 subclades III and V recovered from the mesopelagic (25). Our genomes have generally higher GC content and much lower expected genome sizes than those predicted by Landry *et al*., although these calculations are likely complicated by the relative incompleteness of their genomes (8-47%). The Landry *et al*. genomes indicated a role for SAR202 in the oxidation of recalcitrant dissolved organic matter, and specifically cyclic alkanes, via flavin mononucleotide monooxygenases (FMNOs) and different dehydrogenases that occurred in paralogous groups (25). We observed many of the same gene expansions, namely that of MFS transporters and short-chain dehydrogenases (and related genes), but we did not recover any FMNOs of SFams 4832 or 4965, suggesting subclade and/or niche-specific adaptations. Furthermore, we observed *napAB* and *nosZ* genes for nitrate and nitrous oxide reduction (and expression of these genes), which were not reported for subclade III or V. Our nGOM hypoxia SAR202 genomes had CAZy genes implicating them in degradation of complex compounds such as chitin and pectin. The emerging picture of these taxa from both shallow hypoxic waters and the mesopelagic is one of recalcitrant carbon degraders, with overlapping suites of paralogous genes, but that may be specialized for specific compounds more commonly available in their respective habitats.

This study has also developed roles for CP taxa in a shallow marine water column during hypoxia. The most active organism in our survey based on the ratio of RNA to DNA reads recruited, Bin 13, putatively belongs to a group with little genomic data-ACD39. The original ACD39 genome was reconstructed from an aquifer community (33). Although this was only a partial genome, it shared some features with our putative ACD39 member, namely pilin and chemotaxis genes, those containing TPR and tetratricopeptide repeats, and CAZy genes for degradation of complex compounds such as starch (33). Our study provides evidence that these taxa have relatively large genomes (~4.8 Mbp), are active aerobes in nGOM hypoxia, and have chemotaxis and motility genes that could facilitate scavenging and surface attachment. However, most of the highly expressed genes in this organism were annotated as hypothetical proteins, so much of the function of these organisms remains to be uncovered.

Bins 50 and 48 provide novel genome data for bacterioplankton in nGOM hypoxia, although the exact taxonomic position of these bins remains in conflict. The ribosomal protein tree provides evidence that these taxa belong to the Latescibacteria (WS3) (Fig. S1), but 16S rRNA genes (Fig. S6) and our amplicon data point toward membership in the more poorly understood PAUC34f clade. Since no previous genome data exist for PAUC34f, we cannot rule out erroneous assignment in the ribosomal protein tree due to insufficient taxon selection. Bins 50 and 48 represented the largest genomes of the study, with estimated complete sizes of ~5-6 Mbp, and numerous genes suggesting degradation of a wider suite of complex organic matter than any of the other genomes examined. For example, they were the only genomes with numerous polysaccharide lyase genes, and these likely facilitate breakdown of algal cell wall components like pectin and alginate. The Bin 50 genome was among the most active across all samples (Fig. 3), and we detected expression of cytochrome c oxidase genes, and those for DNRA, in both the Bin 50 and 48 genomes. Thus, we expect these organisms to have an aerobic, potentially facultatively anaerobic, multifaceted chemoorganoheterotrophic metabolism with roles in complex carbon compound degradation (like that of algal cell walls) and the nitrogen cycle.

If these bins belong to PAUC34f, they represent the first genomic data for the group. Although originally discovered, and commonly found, in marine sponges (32, 62-64), this putative bacterial phylum (via GreenGenes/SILVA) has been detected as a rare group in other marine invertebrates (65) and stream sediment (66), and we identified 18 distinct but rare PAUC34f OTUs in nGOM hypoxia, compared to just three from WS3 (21). Although the majority of studies suggest an endosymbiotic lifestyle for PAUC34f, our representative genome data point towards a free-living existence with multiple terminal electron accepting processes, motility genes for seeking more favorable conditions, and a large metabolic repertoire for degradation of complex compounds. On the other hand, if these genomes represent WS3, the sister clade to PAUC34f (Fig. S6), they have many similarities to the lifestyles inferred from recent metagenomic investigations (67, 68). Specifically, while this group was previously considered anaerobic (67), new data have supported an aerobic lifestyle for some members (48), and revealed complete electron transport chains and both high and low affinity cytochrome c oxidases (68). The Bin 50 and 48 genomes predict aerobic metabolism as well, although only with low affinity cytochrome c oxidases. Farag *et al*. also found little evidence of these taxa in host-associated environments, contrary to PAUC34f sequence data (68). The enrichment of PL family genes in Bins 50 and 48, polysaccharide degradation capability in general, and specific genes for degradation of cell wall components, all corroborate previous findings on WS3 as well (68). Bin 50 had 78 annotated peptidases, nearly double that in all other genomes in the study (Bin 48 had 46), which also concurs with metagenomic predictions for WS3 (68). Our genomes differed from WS3 metagenomes principally in the predicted DNRA metabolism and the dramatic expansion of sulfatases. Although sulfatases were observed in WS3 metagenomes (68), they were not present in the numbers associated with Bin 50 (n=102). A large cadre of sulfatases has been previously reported for *Lentisphaera* (n=267) and *Pirellula* (n=110) genomes (69, 70) and suggests specialization for degradation of sulfate-esters to satisfy carbon and/or sulfur requirements.

Although Parcubacteria and Peregrinibacteria occurred in low abundance (Fig. S7) and we detected activity in only a few samples, their recovery in the hypoxic zone is notable because these organisms have generally been associated with anoxic environments. Our predicted genome sizes (~1.5 Mbp) corroborate previous reports of these organisms having small genomes (31, 48). We did not observe any genes associated with nitrogen or sulfur redox transitions, although we cannot rule these capabilities entirely due to incomplete genomes. Regardless, we can hypothesize that Parcubacteria and Peregrinibacteria persist as members of the rare biosphere until they can take advantage of microanoxic niches in the water column where they participate in carbon cycling as obligately fermentative organisms.

Excluding Parcubacteria and Peregrinibacteria, the other uncultivated groups in the nGOM hypoxic zone had one or more genomes that encoded cytochrome c oxidases (and other electron transport chain components) for respiring oxygen, making these taxa likely only facultative anaerobes. Pervasive aerobic metabolism in an oxygen-depleted water column may seem counterintuitive, yet despite DO being as low as 2.2 μmol kg^-1^ in the E2A sample, oxygen probably remained high enough to sustain aerobic microbes. As little as ~0.3 μmol kg^-1^ oxygen inhibited denitrification in OMZ populations by 50% (71), and even *Eschericia coli* K-12 could grow aerobically at oxygen concentrations as low as 3 nM (72). Thus, for many organisms, active aerobic respiration likely persists even in suboxic waters during nGOM hypoxia.

Nevertheless, our data also suggests pervasive co-reduction of alternative terminal electron acceptors (oxygen, nitrate, nitrite, nitrous oxide, and sulfur), sometimes within the same organism (Fig. 4). Co-reduction of electron acceptors with different redox potentials across a community could indicate microniches and/or aggregates in the water column where DO concentrations drop below bulk values (40). Alternatively this can occur with an abundance of electron donor, and overlapping redox processes have been reported in multiple environments, including aquatic ones (50, 73). Concurrent expression of genes for multiple terminal electron accepting processes within a single organism has been proposed as a means of improved readiness for dynamic conditions, albeit at the cost of lower productivity (40). Given that many uncultivated taxa likely perform multiple terminal electron accepting processes (and possibly do so simultaneously), and we found a comparative cornucopia of genes for degradation of chemoorganohetrotrophic energy sources, we hypothesize that niche differentiation within uncultivated hypoxic zone bacterioplankton occurs predominantly via specialization for different oxidizable substrates rather than for distinct roles in the canonical redox cascade (4, 5).

Importantly, many of the active uncultivated taxa also appeared adapted for degradation of complex carbon substrates. Such compounds might comprise the bulk of available organic matter during the later stages of hypoxia after initial oxygen depletion by microorganisms feeding on more labile carbon sources. Selection for chemoorganotrophic microbes adapted to utilize recalcitrant organic matter could also explain why organisms that do not require an exogenous carbon source, such as the chemolithoautotrophic *Nitrosopumilus*, proliferate during hypoxia (21, 22) compared to their levels during spring before DO decreases (74, 75). Temporal data on the relative abundance and activity of these nGOM microbial dark matter organisms, and of organic matter composition in the water column, will be critical to more fully understand the relationship of bacterioplankton to the creation, maintenance, and dissolution of nGOM hypoxia.

## Materials and Methods

### Sample selection and nucleic acid processing

Six samples representing hypoxic (n=4) and oxic (n=2) DO concentrations were picked from among those previously reported (21) at stations D1, D2, D3, E2, E2A, and E4 (Table S1). DO, and nutrient collection information is detailed in Gillies *et al*., 2015. Nucleic acids were collected as follows: At these six stations 10 L of seawater was collected and filtered with a peristaltic pump. A 2.7 μM Whatman GF/D pre-filter was used and samples were concentrated on 0.22 μM Sterivex filters (EMD Millipore). Sterivex filters were immediately sparged, filled with RNAlater, and placed at −20°C, at which they were maintained until extraction. DNA and RNA were extracted directly off of the filter by placing half of the Sterivex filter in a Lysing matrix E (LME) glass/zirconia/silica beads Tube (MP Biomedicals, Santa Ana, CA) using the protocol described in Gillies *et al*. (2015) which combines phenol:chloroform:isoamyalcohol (25:24:1) and bead beating. Genomic DNA and RNA were stored at −80°C until purified. DNA and RNA were purified using QIAGEN (Valencia, CA) AllPrep DNA/RNA Kit. DNA quantity was determined using a Qubit2.0 Fluorometer (Life Technologies, Grand Island, NY). RNA with an RNA integrity number (RIN) (16S/23S rRNA ratio determined with the Agilent TapeStation) ≥ 8 (on a scale of 1-10, with 1 being degraded and 10 being undegraded RNA) was selected for metatranscriptomic sequencing. Using a Ribo-Zero kit (Illumina) rRNA was subtracted from total RNA. Subsequently, mRNA was reverse transcribed to cDNA as described in Mason *et al*. (2012) (76).

### Sequencing, assembly, and binning

DNA and RNA were sequenced separately, six samples per lane, with Illumina HiSeq 2000 chemistry to generate 100 bp, paired-end reads (180 bp insert size) at the Argonne National Laboratory Next Generation Sequencing facility. The data are available at the NCBI SRA repository under the BioSample accession numbers SAMN05791315-SAMN05791320 (DNA) and SAMN05791321-SAMN05791326 (RNA). DNA sequencing resulted in a total of 416,924,120 reads that were quality trimmed to 413,094,662 reads after adaptors were removed using Scythe (https://github.com/vsbuffalo/scythe), and low-quality reads (Q < 30) were trimmed with Sickle (https://github.com/najoshi/sickle). Reads with three or more Ns or with average quality score of less than Q20 and a length < 50 bps were removed. Genomes were reconstructed using two rounds of assembly. Metagenomic reads from all six samples were pooled, assembled, and binned using previously described methods (77, 78). Briefly, quality filtered reads were assembled with IDBA-UD (79) on a 1TB RAM, 40-core node at the LSU High Performance Computing cluster SuperMikeII, using the following settings: ‐mink 65 –maxk 105 –step 10 – pre_correction –seed_kmer 55. Initial binning of the assembled fragments was performed using tetra-nucleotide frequency signatures using 5 kbp fragments of the contigs. Emergent self-organizing maps (ESOM) were manually delineated and curated based on clusters within the map. The primary assembly utilized all reads and produced 28,080 contigs ≥ 3 kb totaling 217,715,956 bp. Of these, 303 contigs were over 50 kb, 72 over 100 kb, and the largest contig was just under 495 kb. Binning produced 76 genomes, of which 20 genomes were assigned to lineages with uncultivated representatives using CheckM, ribosomal protein trees, and 16S rRNA gene sequences (below).

### DNA and RNA mapping

Metagenomic and metatranscriptomic sequencing reads from each sample were separately mapped to binned contigs using BWA (80) to compare bin abundance across samples and facilitate bin cleanup (below). Contigs within each bin were concatenated into a single fasta sequence and BWA was used to map the reads from each sample to all bins. All commands used for these steps are available in supplementary information.

### Bin QC

Bins were examined for contamination and completeness with CheckM (35), and we attempted to clean bins with > 10% estimated contamination using a combination of methods. First, the CheckM modify command removed contigs determined to be outliers by GC content, coding density, and tetranucleotide frequency. Next, in bins that still showed > 10% contamination, contigs were separated according to comparative relative abundance of mean DNA read coverage by sample. Final bins were evaluated with CheckM again to generate the statistics in Table S1 and final bin placements in the CheckM concatenated gene tree (Fig. S2).

### Ribosomal protein tree

The concatenated ribosomal protein tree was generated using 16 syntenic genes that have been shown to undergo limited lateral gene transfer (rpL2, 3, 4, 5, 6, 14, 15, 16, 18, 22, 24 and rpS3, 8, 10, 17, 19) (81). Ribosomal proteins for each bin were identified with Phylosift (82). Amino acid alignments of the individual ribosomal proteins were generated using MUSCLE (83) and trimmed using BMGE (84) (with the following settings: -m BLOSUM30 –g 0.5). The curated alignments were then concatenated for phylogenetic analyses and phylogeny inferred via RAxML v 8.2.8 (85) with 100 bootstrap runs (with the following settings: mpirun -np 4 -npernode 1 raxmlHPC-HYBRID-AVX -f a -m PROTCATLG -T 16 -p 12345 -x 12345 -# 100). Note this is similar to the number utilized in a previous publication for this tree with automated bootstrapping (86), and required just over 56 hours of wall clock time. The alignment is available in SI.

### Average amino acid identity

AAI was calculated with Get Homologues (87) v. 02032017, with the following settings: –M –t 0 –n 16 –A.

### Taxonomic assignment

Taxonomy for each bin was assigned primarily using the ribosomal protein tree. However, for bins that did not have enough ribosomal proteins to be included in the tree, or for which the placement within the tree was poorly supported, assignments were made based on the concatenated marker gene tree as part of the CheckM analysis (Fig. S2), or via 16S rRNA gene sequences, when available. 16S rRNA genes were identified via CheckM, and these sequences were aligned against the NCBI nr database using BLASTN to corroborate CheckM assignments. In the case of the SAR202 genomes, which did not have representative genomes in either the ribosomal protein tree or the CheckM tree, the 16S rRNA gene sequences for two of the three bins (43-1, 43-2) were available and aligned with the sequences used to define the SAR202 clade (24) (Fig. S5). Alignment, culling, and inference were completed with MUSCLE (83), Gblocks (88), and FastTree2 (89), respectively, with the FT_pipe script. The script is provided in SI. The 16S rRNA gene tree for subclade assignment of SAR406 (Fig. S4) was assembled by blasting the four 16S sequences predicted by CheckM against a local GenBank nt database using blastn (v. 2.2.28+) (90), selected the top 100 non-redundant hits to each sequence, and manually removing all hits to genome sequences. These were combined with previously defined SAR406 subclade reference sequences (26), fosmid 16S sequences (27), single-cell genome sequences (23), and run through alignment, culling, and inference with FT_pipe. Taxa with identical alignments were removed with RAxML v 8.2.8 (85) using default settings, and the final tree was inferred using FastTree2 (89). For putative CP genomes, taxonomy was also evaluated by examining the taxonomic identification for each of the predicted protein sequences after a BLASTP search against the NCBI nr database. Post-blast, the number of assignments to the dominant one or two taxonomic names, along with the number of assignments to “uncultured bacterium,” was plotted for each genome according to the bit score quartile (Fig. S3). Quartiles were determined in R using the summary function. Bin 56 has two ribosomal protein operons on scaffold_2719/Ga0113622_1153 and scaffold_21777/Ga0113622_1009. In the ribosomal protein tree, the former placed the organism in the *Planctomycetaceae*, while the latter (which was much smaller) placed the organism in CP WS3. The majority of BLASTP annotations to the nr database matched *Planctomycetaceae* taxa, as did the 16S rRNA gene sequences found in the genome, so Bin 56 organism was designated a Planctomycetes and not WS3, and excluded from this study. The 16S rRNA gene from Bin 50 was also used to infer taxonomic identity using and established phylogeny for the WS3 clade (68) and relevant outgroups. The Bin 50 sequence was blasted against the greengenes database (Dec. 2013) with megablast, and since many of the top hits belonged to the PAUC34f clade, these were included with the sequences from Farag *et al*. 2017. Alignment, culling, and inference was completed with FT_pipe. Node labels were constructed with the newick utilities (91) script nw_rename.

### Metabolic reconstruction

Post-binning, genomes were submitted individually to IMG (92) for annotation. Genome accession numbers are in Table S1, and all are publically available. Metabolic reconstruction found in Table S1 and Figs S5-7/ S11-13, came from these annotations and inspection with IMG’s analysis tools, including KEGG pathway assignments and transporter predictions. Transporters highlighted for DOM uptake were identified based on information at the Transporter Classification Database (93). Carbohydrate-active enzymes (CAZymes) were predicted using the same routines as those in operation for the updates of the carbohydrate-active enzymes database (www.cazy.org) (94).

### RPKM abundance of taxa and genes

Abundance of taxa within the sample was quantified by evaluating mapped reads using Reads-Per-Kilobase-Per-Million (RPKM) normalization (95) according to *A*_*ij*_ = (*N*_*ij*_/*L*_*i*_) x (1/*T*_*j*_), where *A*_*ij*_ is the abundance of bin *i* in sample *j*, *N*_*ij*_ is the number of reads that map to bin from sample *j*, *L*_*i*_ is the length of bin *i* in kilobases, and *T*_*j*_ is the total number of reads in sample *j* divided by 10^6^. These values were generated for all bins, with only the data for the 20 uncultivated bins reported here. All contigs within a given bin were artificially concatenated into “supercontigs” prior to mapping. *N*_*ij*_ was generated using the samtools (80) idxstats function after mapping with BWA. The data in Fig. S7 were created by summing (*N*_*ij*_/*L*_*i*_) for groups of taxa defined in Table S1 prior to multiplying by (1/*T*_*j*_). RNA coverage was used to evaluate both bin and gene activity for all bins. Mean coverage for each supercontig was calculated using bedtools (96) and bins were assigned a rank from lowest mean recruitment (1) to highest mean recruitment (2). Bins with particularly high or low activity (transcript abundance) relative to their abundance (genome abundance) were identified using rank-residuals, calculated as follows: On a plot of DNA coverage rank vs. RNA coverage rank, residuals for each bin or gene were calculated from the identity. As the rank-residuals followed a Gaussian distribution, bins with a residual that was > 1 s.d. from the rank-residual mean were classified as having higher-than-expected transcriptional activity; bins with a residual that was < 1 s.d. from the mean were classified as having lower-than-expected transcriptional activity. RPKM values were also calculated for every gene in every bin analogously to that for bins, using RNA mapping values extracted with the bedtools multicov function. Sample E2 was omitted from gene-specific calculations as only 4588 transcriptomic reads mapped successfully from this sample, compared to >100,000 from other samples. 17,827 of 140,347 genes had no evidence of expression in any sample and so were removed from further analysis. 3,840 genes recruited reads in all remaining samples. All calculations are available in Table S1 or the R markdown document Per.gene.RPKM.Rmd in Supplemental Information. Table S1 includes only analyzed data for the uncultivated bins reported in this study. Note that RPKM values indicate abundance measurements across a small number of samples. While we can evaluate the relative expression of genes for those samples, our dataset lacks sufficient power to evaluate estimates of significance in differential expression.

### nrfA sequence assessment

Initial annotation of our bins identified putative homologs to the *nrfAH* genes associated with dissimilatory nitrite reduction to ammonia. Since *nrfA-type* nitrite reductases can be misannotated due to homology with other nitrite reductases, annotation for these genes was curated with phylogenetic analysis using known *nrfA* genes (38) obtained via Dr. Welsh (personal communication). Alignment, culling, and inference were completed with the FT_pipe script. The tree was rooted on the designated outgroup octaheme nitrite reductase sequence from *Thioalkalivibrio nitratireducens* ONR. Node labels were constructed with the newick utilities (91) script nw_rename. Visualization of the alignment (Fig. S8B,C) to confirm the presence of the first CXXCK/CXXCH and highly conserved KXQH/KXRH catalytic site was completed with the MSAViewer (97) online using the un-culled *nrfA* alignment as input.

### SFam homology searches

To identify group specific expansions in particular gene families, we performed a homology search of all predicted protein coding sequences in each bin against the Sifted Families (SFam) database (41) using hmmsearch (HMMER 3.1b (98)) with default settings except for the utilization of 16 cpus per search.

## Acknowledgements

The authors thank the crew of the R/V *Pelican*, Dr. Allana Welsh and Dr. Mostafa Elshahed for providing fasta files for *nrfA* and WS3 phylogenetic comparisons, and Dr. Elshahed for helpful comments regarding WS3 phylogeny. Portions of this research were conducted with high performance computing resources provided by Louisiana State University (http://www.hpc.lsu.edu).

## Author contributions

JCT and OUM designed the study. LEG, NNR, and JCT collected samples. NNR provided processed oceanographic data. LEG and OUM extracted, quantified, and determined quality of nucleic acids. JCT, KWS, and BJB reconstructed the genomes. KWS, BJB, BT, BH, and JCT conducted downstream analyses. JCT led manuscript writing and all co-authors evaluated and contributed edits.

## Supplemental Information

**Supplemental Text** provides additional information on taxonomic assignments.

**Table S1.** Spreadsheet (Table_S1.xlsx) containing information on taxonomy, CheckM results, IMG statistics, partial metabolic reconstruction, gene annotations associated with Figures 1 and 2, transporter classifications, CAZy predictions, sample chemical data, RPKM values and gene neighborhoods for WS3 cytochrome c oxidases and *nrfA* genes from SAR406 and WS3.

**Figure S1.** Maximum likelihood tree of concatenated ribosomal protein coding genes. Values at internal nodes indicate bootstrap support (n=100). Scale bar indicates changes per position.

**Figure S2.** Phylogenetic placement of bins based on CheckM.

**Figure S3.** Annotations of protein-coding gene sequence best blastp hits in the nr database, divided into quartiles by bit score, for CP bacteria.

**Figure S4.** SAR406 16S rRNA gene phylogeny. Genes recovered from assembled bins have 45* or 51* designations. Values at nodes indicate Shimodaira-Hasegawa “like” values (89). Scale bar indicates changes per position.

**Figure S5.** SAR202 16S rRNA gene phylogeny. Genes recovered from assembled bins are indicated as 43-*. Subclades are designated according to Morris *et al*. 2004, and the tree is rooted according to Figure 1 in that publication. Values at nodes indicate Shimodaira-Hasegawa “like” values (89). Scale bar indicates changes per position.

**Figure S6.** 16S rRNA gene phylogeny of the WS3 clade (68) with added PAUC34f sequences from the GreenGenes database and the Bin 50 sequence. Tree is rooted on the Archaea according to Fig. S1 in Farag *et al*. 2017. Values at nodes indicate Shimodaira-Hasegawa “like” values (89). Scale bar indicates changes per position.

**Figure S7.** Metagenomic RPKM values for each group, comprised of aggregated values for each bin within the group. Values are plotted according to sample and colored according to the dissolved oxygen (DO) concentration from where the sample was taken.

**Figure S8.** Evaluation of predicted nrfA genes in SAR406 and Bins 50/48. A) Phylogenetic tree of predicted *nrfA* genes. Additional taxa are from Figure 3 in Welsh *et al*., 2014. The tree was rooted at the midpoint. Values at nodes indicate Shimodaira-Hasegawa “like” values (89). Scale bar indicates changes per position. B&C) Conserved catalytic motifs within the *nrfA* gene. B) Black square surrounds the first heme-binding CXXCK/CXXCH motif. C) Black square surrounds the catalytic KXQH/KXRH motif. The alignment follows highlighting found in Welsh *et al*., 2014 (38). All genes numbers from this study are indicated as 26536*, corresponding to rows 64 and 66-68.

## Additional Supplemental Information

Such as scripts, workflows, and key files, including fasta files for each tree, are provided as a link hosted at the Thrash Lab website: http://thethrashlab.com/publications.

